# Apoptosis is increased in cortical neurons of female Marfan Syndrome mice

**DOI:** 10.1101/2024.10.14.618278

**Authors:** Mitra Esfandiarei, Faizan Anwar, Manogna Nuthi, Alisha Harrison, Mary Eunice Barrameda, Tala Curry, Kasey Pull, Theresa Currier Thomas, Nafisa M. Jadavji

## Abstract

Marfan Syndrome (MFS) is an autosomal dominant genetic disorder that affects connective tissue throughout the body due to mutations in the *FBN1* gene. Individuals with MFS display symptoms in different organs, particularly in the vasculature, but the mechanisms of this multi-system dysfunction are still under investigation. There is still a gap in our understanding of the impact of monogenic connective tissue aberrations on the brain. This study aims to determine the impact of MFS on neurodegeneration, in cortical brain tissue of male and female MFS mice. Brain tissue of 6-month-old female and male mice with the *FBN1*^*C1041G/+*^ mutation and wildtype litter mates was collected and stained for active caspase-3 (ac3), brain derived neurotrophic factor (BDNF), and neuronal nuclei (NeuN) or with TUNEL and DAPI. Data revealed increased levels of ac3 in neurons within the sensory and motor cortical areas of female MFS mice compared to sex- and age-matched controls. We confirm increased levels of apoptosis in MFS mice using TUNEL staining within the same brain areas. We also report increased levels of neuronal BDNF levels in cortical brain tissue of male and female MFS mice. These results indicate a heightened susceptibility for neurodegeneration in the mouse model of MFS.

**Significance Statement:** The study revealed that female mice with a *FBN1*^*C1041G/+*^ mutation have more apoptotic neurons in sensory and motor cortical brain tissue compared to wildtype litter mates.

## Introduction

Marfan syndrome (MFS) is a hereditary systemic disorder of the connective tissue caused by mutations in the gene encoding for the large glycoprotein fibrillin 1 (FBN1) that affects the cardiovascular, pulmonary, musculoskeletal, and ocular systems [1]. MFS can lead to manifestations such as aortic root aneurysm, scoliosis, pneumothorax, and lens dislocation, with aortic root dissection and rupture considered the most life-threatening complications [2].

The development of animal models of MFS has been instrumental in understanding the underlying mechanisms contributing to vascular complications in MFS. The use of *FBN1* mutant mice has shed light on how mutations in the *FBN1* gene can disrupt the regulation of the downstream protein transforming growth factor beta (TGF-β) protein [3]. This disruption results in increased expression and activity of matrix metalloproteinases 2 and 9 (MMP-2 & -9), leading to extracellular matrix degradation and weakening of connective tissues, particularly in vital organs such as the heart, lungs, aorta, and brain [4–6]. In addition, significant increases in tissue proteases MMP-2 and -9 can enhance inflammatory responses, exacerbating the pathology [7].

Despite recent advances in understanding the mechanisms involved in the development of MFS-associated aortopathy, there has been limited focus on the potential neurological and cerebrovascular implications. Previous clinical studies in MFS patients have revealed complications such as impaired cerebral blood flow, epilepsy, cephalgia, and varicose spinal veins, with reported increases in chronic migraine, neurodivergence, and intracranial aneurysms and stroke [8–10]. We have also recently reported a significant decrease in cerebral blood flow (CBF) in the mouse model of MFS (*FBN1* ^*C1041G/+*^) [11], underscoring the potential impact of FBN1 abnormalities on cerebrovascular and brain function.

Reduced CBF is linked to cognitive decline in patients with dementia [12]. Other studies have reported that reduced CBF can lead to hypoxia and nutrient deprivation, triggering oxidative stress, inflammation, cellular stress response, and apoptosis, resulting in neural damage and cell death [13]. Apoptosis, or programmed cell death, is a fundamental process in the brain, playing key roles in maintaining normal brain homeostasis, establishing neural networks, removing damaged cells and proteins, and responding to brain injuries [14]. Dysregulation is such an important and corrective mechanism which can lead to various brain pathologies, including neurodegenerative diseases and impaired recovery from injury [15,16].

In this study, we aim to investigate the presence and extent of apoptosis in the brain tissue of both male and female MFS mice (*FBN1* ^*C1041G/+*^), with the aim of filling the existing knowledge gap regarding brain pathology in MFS. We anticipate that this study provides new insights into the broader implications of this connective tissue disorder, suggesting more effective therapeutic approaches to improve the quality of life in individuals with MFS.

## Materials and Methods

### Experimental Animals

The mouse model used in the study carries a heterozygous missense mutation in the *FBN1* gene (*FBN1*^*C1041G/+*^, cysteine to glycine substitution Cys^1041^→Gly), recapitulating the aortic aneurysm phenotype observed in MFS patients [11,17,18]. A breeding colony was established by crossing male MFS (*FBN1*^*C1041G/+*^) with female control (*FBN1*^*+/+*^) mice obtained from The Jackson Laboratory (Bar Harbor, ME). Male and female mice were housed in the institutional animal facility with standard animal room conditions (25 °C, 12-hour light-dark cycles, ≤5 mice in a cage). All animals used in this study were cared for in compliance with the Guide for the Care and Use of Laboratory Animals and all experimental procedures were performed according to the Midwestern University Institutional Animal Care & Use Committee (IACUC) approved protocol [reference number AZ-2936]. At 6 months of age, female and male MFS and control littermates (n = 4-6/group) were euthanized using 5% isoflurane (confirmation by pedal reflex), followed by cervical dislocation.

### Immunofluorescence of Brain Tissue

Female and male mice brain tissue was removed and fixed in 4% paraformaldehyde (PFA) for 24 hours, cryo-sectioned at 20µM thickness and serially mounted onto slides. Microscope slides were stored at -80°C until immunofluorescence experiments. To investigate neurodegeneration and plasticity, immunofluorescence analysis of brain tissue was performed. To assess apoptosis-associated neurodegeneration cortical brain sections were stained with active capase-3 primary antibodies (ac3, 1:100, Cell Signaling Technologies, catalog: #9662). All brain sections were also stained with a marker for neuronal nuclei (NeuN, 1:200, Abcam, catalog # ab177487). We measured plasticity using brain-derived neurotrophic factor (BDNF, 1:100, Abcam, catalog #: ab108319). Brain sections were incubated overnight at 4°C with primary antibodies diluted in 0.5% Triton X, and with secondary antibodies Alexa Fluor 488 or 555 (1:200, Cell Signaling Technologies, catalog# 4408 and 4413) at room temperature for 2 hours, followed by staining with 4’, 6-diamidino-2 phenylindole (DAPI, 1:10000, Fisher Scientific, catalog # EN62248). Microscope slides were cover slipped with Fluro-mount mounting media (Fisher Scientific, catalog # OB100-01) and stored at 4°C until imaging.

### Imaging & Statistical Analysis

Stained cortical brain sections (4 to 6 per animal) were imaged using a Leica TCS SPE confocal microscope to create z-stacks. Two observers, who were blinded to experimental conditions, completed cell count colocalization of ac3 and NeuN and TUNEL analysis using ImageJ (NIH). GraphPad Prism 10.02 was used to analyze immunofluorescence staining measurements.

In the GraphPad Prism, D’Agostino-Pearson normality test was performed prior to two-way ANOVA analysis, when comparing the mean measurement of both sex and genotype groups for immunofluorescence staining. Significant main effects of two-way ANOVAs were followed up with Tukey’s post-hoc test to adjust for multiple comparisons. All data are presented as mean ± Standard Error of the Mean (SEM). Statistical tests were performed using a significant level of *p* ≤ 0.05. Brain microscopic images were analyzed by two independent observers who were blinded to experimental treatment groups.

### Results

We assessed the levels of ac3-positive neuronal cells within the cortical tissue of male and female control and MFS mice. **Figure 1A** shows representative images of ac3 and NeuN staining in female control and MFS mice. Our quantification showed an interaction between sex and genotype (**Figure 1B**, F (_1, 12_) = 10.38, *p* = 0.0073). There were increased levels of ac3 and NeuN positive cells in female MFS mice brain (*p* < 0.02), with no changes observed in age-matched male MFS mice. There was no effect of sex (*p* = 0.97) or genotype (*p* = 0.121).

**Figure 1.**
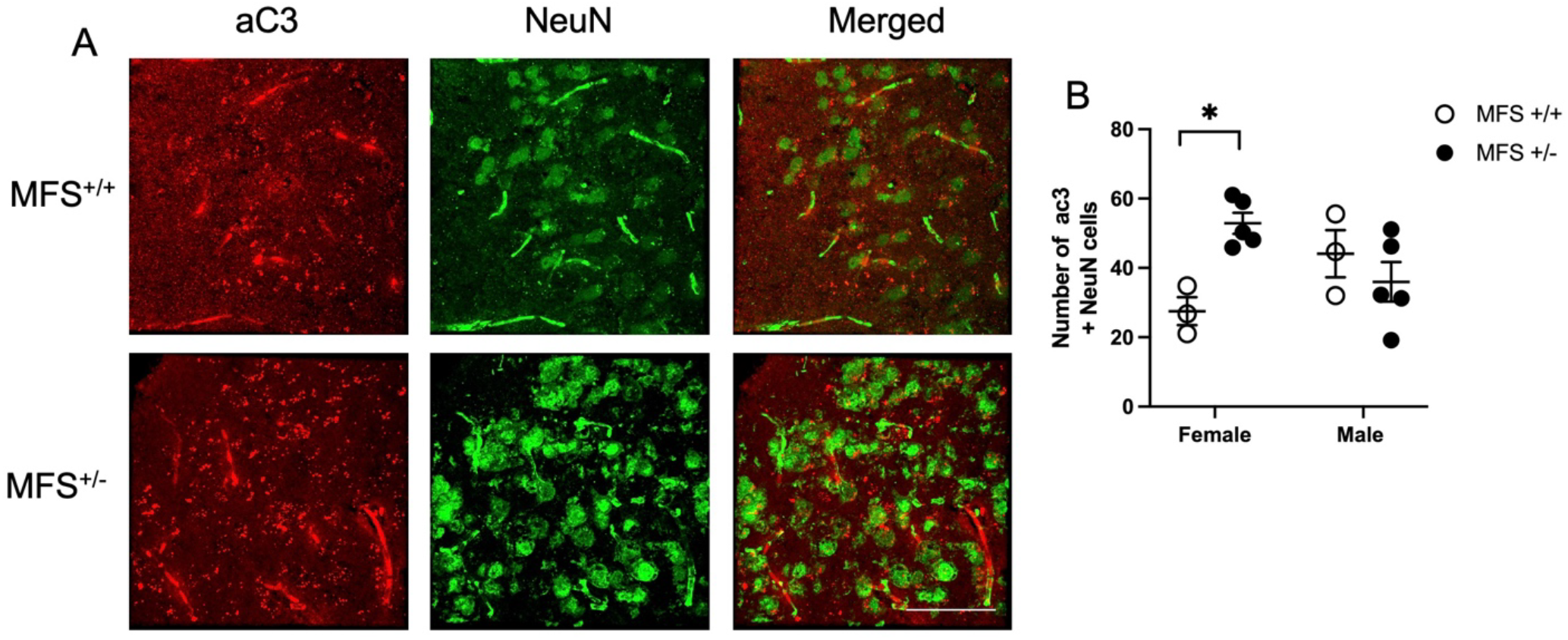
Measurements of active caspase-3 (ac3) and neuron nuclei (NeuN) in motor and sensory motor areas in male and female control and MFS mice. Representative images from female control and MFS mice **(A)** and quantification of ac3 and neuronal nuclei (NeuN) positive cells in brain cross sections from male and female control and MFS mice **(B)**.There was an increase in apoptotic neurons in MFS compared to age- and sex-matched controls (* *p* <0.05, Tukey’s pairwise comparison).

To further confirm apoptosis, we measured levels of TUNEL and DAPI colocalization in cortical brain tissue (**Figure 2A**). There was an increase in MFS mouse brain tissue compared to wildtype controls (**Figure 2B**, F (_1, 8_) = 12.13, *p* = 0.0083). There was no significant interaction (*p* = 0.79) or sex difference (*p* = 0.06).

**Figure 2.**
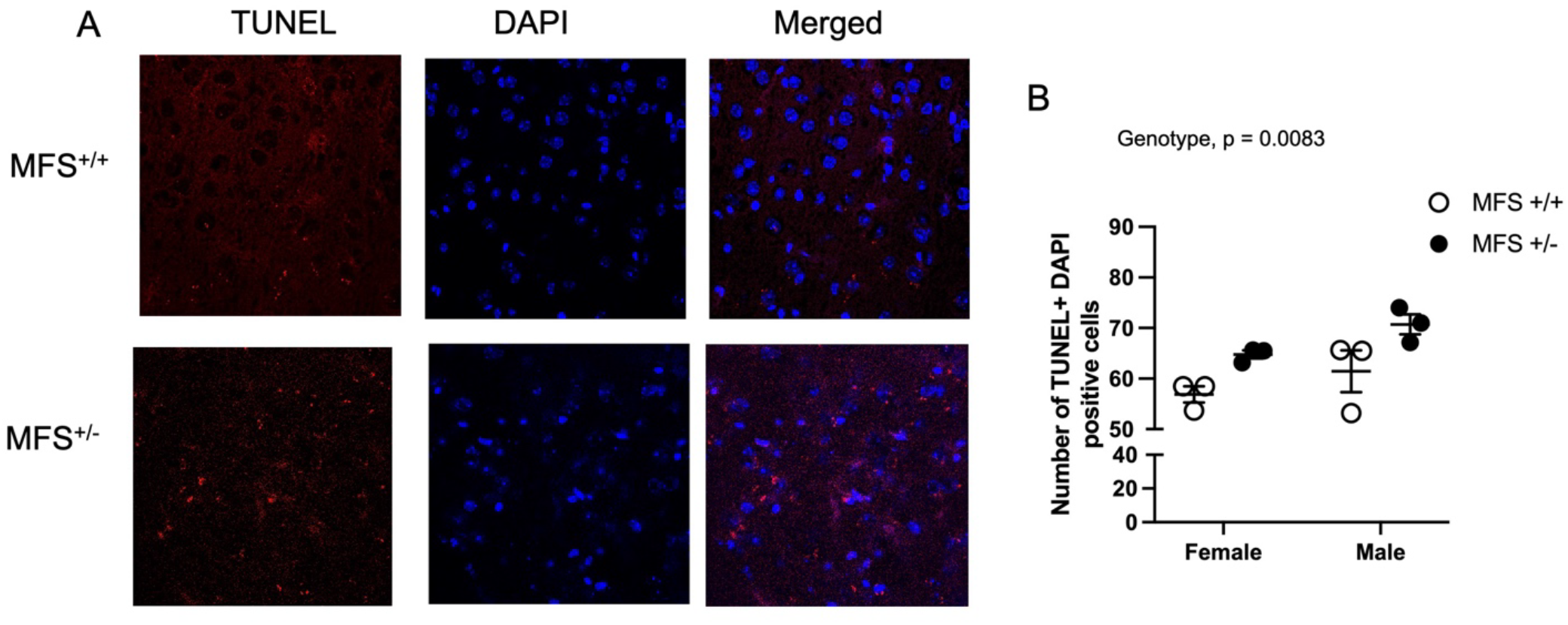
Assessment of TUNEL positive areas in motor and sensory motor areas in male and female control and MFS mice. Representative images from female control and MFS mice **(A)**, and quantification of TUNEL and ′,6-diamidino-2-phenylindole (DAPI) positive cells in brain cross sections from male and female control and MFS mice **(B)**.There was an increase in TUNEL positive cells in female and male MFS compared to controls (*p* = 0.0083).

To measure plasticity within cortical tissue we stained collected mice brain tissue for the brain derived neurotrophic factor (BDNF). Representative images from female MFS mice are shown in **Figure 3A**. As presented in **Figure 3B**, there were increased levels of neuronal BDNF levels in brain tissue in both female and male MFS brain tissue compared to age-matched control littermates (**Figure 3B**, F (_1, 11_) = 8.782, p = 0.012). There were no differences between sex (p = 0.225) or interactions between sex and genotype (p = 0.922).

**Figure 3.**
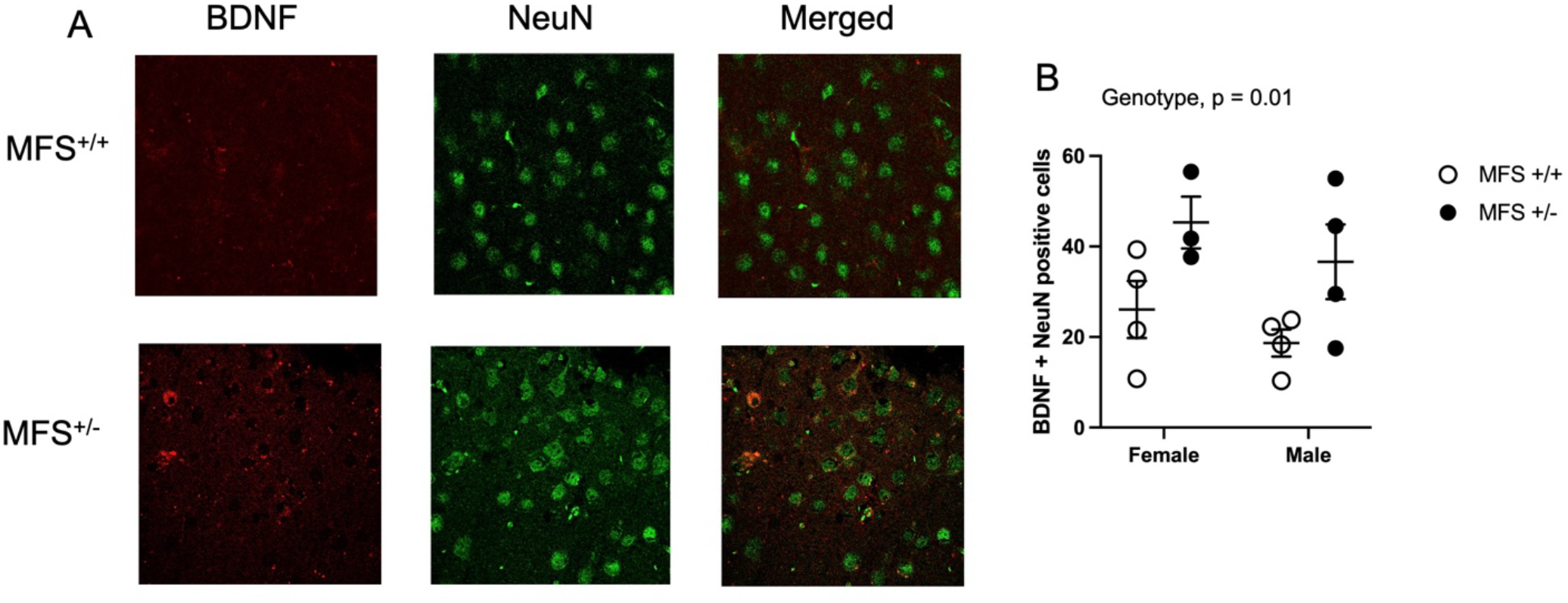
Measurements of brain derived neurotrophic factor (BDNF) and neuron nuclei (NeuN) expression in motor and sensory motor areas in male and female control and MFS mice. Representative images of female mice **(A)** and quantification of BDNF and neuronal nuclei (NeuN) positive cells in both male and female control and MFS mice **(B)**.There was an increase in BDNF neurons in female and male MFS compared to age- and sex-matched control animals (*p* = 0.013).

## Discussion

MFS is primarily known for its impact on cardiovascular, skeletal, and ocular systems. However, evidence suggests that MFS may also involve neurological complications, potentially due to weakened connective tissue in the brain’s vasculature and extracellular matrix. This could lead to structural brain abnormalities, microvascular dysfunction, and even increased vulnerability to neurodegeneration. Investigating cell death mechanisms, such as apoptosis and necrosis, within brain tissue can provide insight into how these processes contribute to neurological symptoms like headaches, migraines, or even cognitive impairments reported in MFS. This study aimed to fill the gap in our understanding of the impact of monogenic connective tissue aberrations on the brain, specially focusing on neurodegeneration of cortical brain tissue in male and female MFS mice. Our study revealed increased levels of active (cleaved) caspase 3 in neurons within the sensory and motor cortical areas of female MFS mice compared to sex-and age-matched controls. We further confirmed increased levels of apoptosis in MFS mice using TUNEL staining within the same brain areas. We also report increased levels of neuronal BDNF levels in cortical brain tissue of male and female MFS mice These results indicate a heightened susceptibility for neurodegeneration in MFS^+/-^ mice, where BDNF may be upregulated in compensation for ongoing cell death or to increase neurotropic support to neurons.

Clinical studies have reported that individuals with MFS have increased risk for cerebrovascular alterations due to their vascular tortuosity phenotype [19]. A recent study investigating the neuropathology in the same MFS mouse model showed a systemic pre-mature aging phenotype is present in the six-month old mice [20]. Specifically, the researchers reported reductions in blood flow of the posterior cerebral artery, as well as increased blood-brain permeability and neuroinflammation [20]. The present study has added to the neurological phenotype of the MFS. Within the sensory and motor cortex, MFS mice showed increased levels of apoptotic neurons compared to wildtype animals along with increased levels of plasticity..

Interestingly, neuronal active caspase-3 levels were significantly increased in the females compared to the males, this is an interesting sex-dependent phenotype in the mouse model of MFS. Clinically, mortality as a result of MFS affects males and females at an equal rate [21]. However, there are sex differences in described outcomes both in clinical and preclinical studies [22]. A possible explanation for the observed sex difference in our present study might be the age of the mice, estrogen has been shown to be neuroprotective in brain response to stress [23,24].

In conclusion, our findings reveal an increased vulnerability to neurodegeneration in the cortical tissue of both male and female MFS mice at 6 months of age. This study is the first to characterize the impact of MFS on neurodegenerative processes in a well-established and commonly used MFS mouse model. Notable, we observed elevated levels of neural plasticity in sexes, which may be associated with the increased levels of neurodegeneration. These novel insights into the neural pathology of MFS underscore the need for further research to delineate the temporal progression of neurodegeneration and to explore additional pathological markers involved in programmed cell death and neural plasticity pathways in the mouse model of MFS.

## Abbreviations

ac3,: active caspase-3:
BDNF: brain derived neurotrophic factor:
FBN1,: large glycoprotein fibrillin 1:
MFS: Marfan syndrome:
NeuN: neuronal nuclei:

## CrediT authorship contribution statement

Mitra Esfandiarei: Conceptualization, Data curation, Formal analysis, Funding acquisition, Investigation, Writing – original draft, Writing – review & editing. Faizan Anwar: Investigation, Writing – review & editing. Manogna Nuthi: Investigation, Writing – review & editing. Alisha Harrison: Investigation, Writing – review & editing. Mary Eunice: Methodology, Validation, Visualization, Writing – review & editing. Tala Curry-Koski: Methodology, Validation, Writing – review & editing. Kasey Pull: Methodology, Validation, Visualization, Writing – review & editing. Theresa Currier Thomas: Conceptualization, Data curation, Formal analysis, Funding acquisition, Investigation, Methodology, Validation, Visualization, Writing – review & editing. Nafisa M. Jadavji: Conceptualization, Data curation, Formal analysis, Funding acquisition, Investigation, Methodology, Validation, Visualization, Writing – original draft, Writing – review & editing.

## Data Availability

Data will be made available upon request.

## Acknowledgements

This research was funded in part by National Institutes of Health, grant number R15HL145646.

